# Differential Analysis of RNA Structure Probing Experiments at Nucleotide Resolution: Uncovering Regulatory Functions of RNA Structure

**DOI:** 10.1101/2021.08.24.457484

**Authors:** Bo Yu, Pan Li, Qiangfeng Cliff Zhang, Lin Hou

## Abstract

RNAs perform their function by forming specific structures, which can change across cellular conditions. Structure probing experiments combined with next generation sequencing technology have enabled transcriptome-wide analysis of RNA secondary structure in various cellular conditions. Differential analysis of structure probing data in different conditions can reveal the RNA structurally variable regions (SVRs), which is important for understanding RNA functions. Here, we propose DiffScan, a computational framework for normalization and differential analysis of structure probing data in high resolution. DiffScan preprocesses structure probing datasets to remove systematic bias, and then scans the transcripts to identify SVRs and adaptively determines their lengths and locations. The proposed approach is compatible with most structure probing platforms (*e*.*g*., icSHAPE, DMS-seq). When evaluated with simulated and benchmark datasets, DiffScan identifies structurally variable regions at nucleotide resolution, with substantial improvement in accuracy compared with existing SVR detection methods. Moreover, the improvement is robust when tested in multiple structure probing platforms. Application of DiffScan in a dataset of multi-subcellular RNA structurome identified multiple regions that form different structures in nucleus and cytoplasm, linking RNA structural variation to regulation of mRNAs encoding mitochondria-associated proteins. This work provides an effective tool for differential analysis of RNA secondary structure, reinforcing the power of structure probing experiments in deciphering the dynamic RNA structurome.

## Introduction

RNA molecules play important roles in myriad cellular processes by forming specific structures^1-3^. Deciphering RNA structure is informative for understanding RNA functions. In recent years, diverse structure probing (SP) methods have been developed to study RNA secondary structure in various cellular contexts, which utilize chemicals that react differentially to nucleotides according to their pairing status^4, 5^. Coupled with next generation sequencing technologies, SP experiments can be performed at high throughput, providing a transcriptome level view of RNA secondary structure^4-6^, *i*.*e*., RNA structurome. There are various mature high-throughput SP platforms, such as SHAPE-Seq^7, 8^, DMS-Seq^9^, and icSHAPE^10^. These platforms offer flexible options for tackling different biological problems, and have achieved success in uncovering pervasive links between RNA structure and RNA function^11^.

Studies have examined how the structures of particular RNA molecules change across multiple cellular conditions, revealing explicit connections between RNA structure and RNA function^12^. For example, the ubiquitous *yybP-ykoY* motif has been shown to adopt distinct structures in response to manganese ions, thus exerting regulatory consequences on protein translation in both *Escherichia coli* and *Bacillus subtilis*^13^. RNA structural variations have also been reported to regulate the binding of trans-acting factors and RNA stability in multiple organisms, including human, yeast, and zebrafish^14-18^.

To explore the SP experiments to uncover dynamic RNA structures, quantitative tools that contrast SP experiments to identify structurally variable regions (SVRs) are in great demand. Several methods have been proposed to identify SVRs. PARCEL^19^ and RASA^20^ directly model and compare raw read counts at each nucleotide position, and then identify regions enriched for position-level signals. However, their models are tailored for specific experimental protocols, and it is not straightforward to extend them to accommodate more emerging SP techniques. StrucDiff^21^, deltaSHAPE^22^, and dStruct^23^ take reactivities as input, which are estimated from raw read counts to summarize the pairing status at each nucleotide position. They search for SVRs with pre-specified search lengths to aggregate differential signals^21-23^. However, the choice of search length is arbitrary.

Despite the success of existing methods, differential analysis of SP data remains challenging in many aspects. First, SVRs manifest great variation in length, ranging from a few to several dozens of nucleotide positions^19, 23, 24^. As a result, searching with fixed search length can lead to insufficient detection power and inaccurate boundary mapping, when the prespecified search length deviates greatly from the true length. Second, SP platforms differ in utilization of additives, specific experimental protocols, and preprocessing pipelines, yielding distinct data types and distributions in output^4, 5, 25^. As a result, it is desirable but usually not easy to extend methods developed for one platform to another. Third, SP data can be confounded by systematic bias^4, 5^, which should be removed from reactivities of the two compared conditions prior to differential analysis^26^. However, to our knowledge, normalization techniques that are compatible with multiple SP platforms are still lacking. Finally, rigorous error control is also highly impactful on the accuracy and biological relevance of output from SVR detection^23^, especially for transcriptome-scale analyses.

To address the unmet needs, we propose DiffScan, a computational framework for differential analysis of SP data at nucleotide resolution. DiffScan normalizes SP reactivities via a built-in Normalization module, which is compatible with various platforms, and then looks for SVRs via a Scan module. The Scan module locates SVRs at nucleotide resolution, and rigorously controls family-wise error rate. We demonstrated with large-scale simulated datasets and benchmark datasets that DiffScan achieves the best power and accuracy in SVR detection across various platforms, compared to state-of-art methods, We applied DiffScan to a recent icSHAPE dataset of RNA structurome in multiple subcellular compartments (e.g., nucleoplasm and cytoplasm), and revealed the potential roles of SVRs in regulating mRNAs encoding mitochondria-associated proteins.

## Results

### Model description

DiffScan comprises a Normalization module and a Scan module (Fig. 1a). SP reactivities of RNA transcripts for two conditions (e.g., *in vitro, in vivo*) are taken as input, and the reactivities are normalized relative to one another in the Normalization module (Fig. 1b) to correct for systematic bias. The corrected reactivities are comparable as far as possible across different cellular conditions. In the Scan module (Fig. 1c), structural variations are initially evaluated at each nucleotide position, and then the algorithm scans through the transcripts with variable search lengths to identify SVRs.

**Fig. 1.**
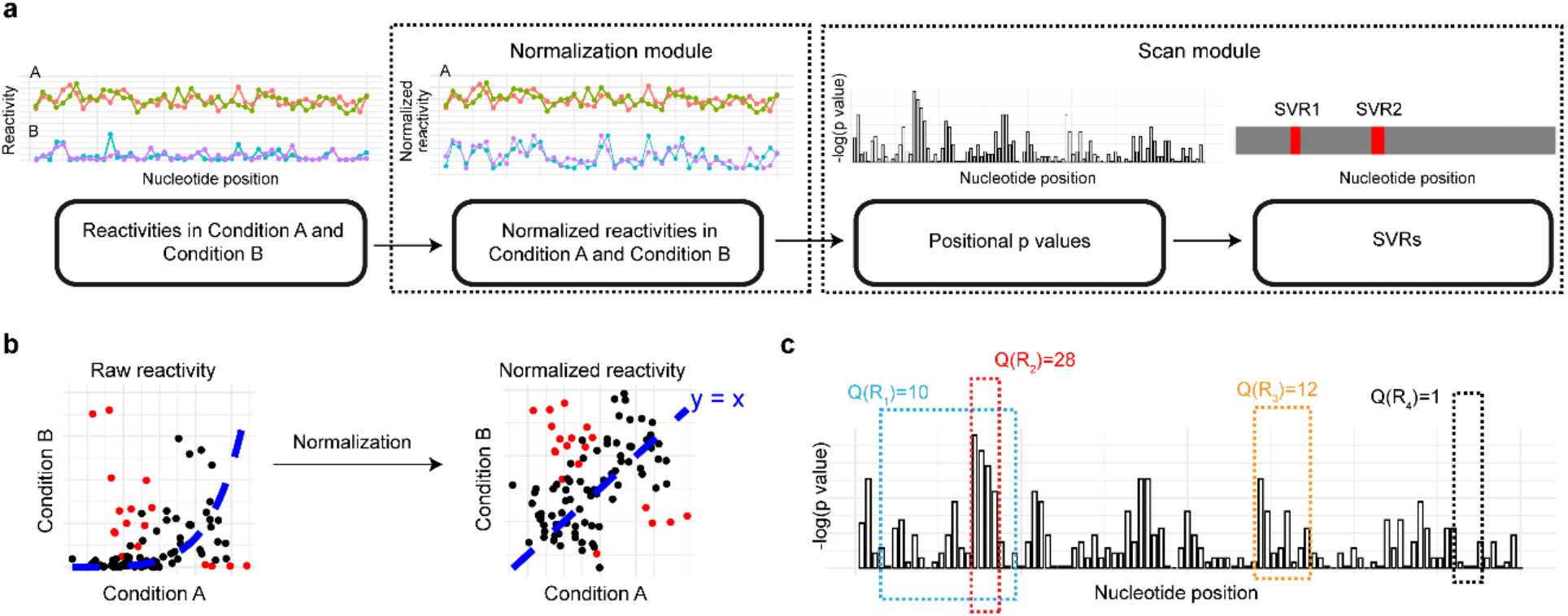
Workflow of DiffScan. **a** Taking raw reactivities as input, DiffScan first normalizes them relative to one another in the Normalization module (b) to correct for systematic bias, and then identifies SVRs in the Scan module (c). Different colors denote reactivity replicates in each condition. **b** The Normalization module transforms raw reactivities into normalized reactivities to remove systematic bias. The normalized reactivities are comparable as far as possible across different cellular conditions. Red points indicate nucleotide positions in SVRs. **c** Taking normalized reactivities as input, the Scan module first calculates the significance of any differential signals for each nucleotide position with Wilcoxon test, and then concatenates positional p values into a regional signal via scan statistic. The significance of the scan statistic for each enumerated region is evaluated by Monto Carlo sampling, and those regions crossing a specified significance threshold are reported as SVRs.

#### Normalization

It is known that the reactivities of particular RNA nucleotide positions can be affected by a variety of confounding factors in SP experiments^26^, including transcript abundance, sequencing depth, and signal-to-noise ratio (see Methods). Owing to discrepancies in these factors, the SP reactivities obtained from two conditions lacking substantial biological differences may show substantial differences^27^ (see Supplementary Fig. 1 for an example). The unwanted variations will then persist throughout the routine preprocessing steps. Thus, in practice careful adjustment between and within conditions is essential when comparing SP data obtained from samples in distinct conditions. Related challenges have been widely reported for other high-throughput sequencing experiments, such as ChIP-Seq^28^ and ATAC-Seq^29^. Inspired by normalization approaches for other high-throughput sequencing based methods, we propose the following normalization procedure for differential analysis of SP data.

To account for a potentially wide range of reactivity levels, we first rescale the reactivity values into the interval [0,1] after a 90% winsorization (see Methods). Next, the reactivities of within-condition replicates, if available, are processed by quantile normalization^30^, so that the quantiles of each replicate are matched within conditions. We subsequently normalize between-condition reactivities using an approach similar to MAnorm^28^. Briefly, a structurally invariant set of nucleotide positions *S* is determined as a “pivot” for normalization, and the reactivities are transformed so that the transformed reactivities are at the same level for the pivot set between conditions. The basic idea is to learn transformation rules from *S* and then extrapolate the learned transformation to all nucleotide positions of the transcripts. In particular, the transformation is learned from training a linear model from *S*, which takes reactivities in one condition as response and reactivities in the other condition as predictor. The adjusted reactivities from the Normalization module are then ready for differential analysis in the Scan module (Supplementary Fig. 2). We show in theory that the transformation explicitly corrects for differences in sequencing depth and signal-to-noise ratio.

#### Scan

Formally, “scan statistics” refer to statistical methods for cluster detection in time and space^31, 32^. These methods can accurately map regional signals and control family-wise error rates during multiple testing of interrelated hypotheses. Scan statistics have been successfully applied to many areas including molecular biology^33^ and human genetics^34^. The Scan module of DiffScan was developed with the goal to identify SVRs at nucleotide resolution in a data-adaptive fashion. In brief, the Scan module slides through the transcripts to enumerate overlapping regions of different lengths, identifies regions with maximal differential signals, and evaluates their statistical significance. In detail, we first quantify positional differential signals by calculating a p value for each nucleotide position by Wilcoxon test, which contrasts reactivities between conditions in a small window surrounding the nucleotide position. Note that the Normalization module does not enforce any specific distributions of the normalized reactivities (Supplementary Fig. 3), and the nonparametric test we use guarantees robust evaluation of differential signals for various SP platforms.

Second, for each region R, we propose the following scan statistic to quantify differential signals for regions,

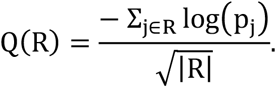

The sum in the numerator aggregates positional differential signals, while the denominator penalizes the extension of a candidate region. Intuitively, regions that are enriched with structurally variable nucleotides will obtain high Q(R) scores.

We search in the transcripts with sliding windows of different lengths, and calculate the scan statistic for each candidate region (Fig. 1c). A Monte Carlo approach is then implemented to evaluate the statistical significance of the scan statistics, which addresses the multiple testing problem for the overlapping regions by controlling the family-wise error rate (See Supplementary Methods). Accordingly, we can identify SVRs with accurately mapped boundaries based on the significance of the scan statistics (Fig. 1c).

### Validation of DiffScan using simulated SP datasets

We simulated RNA secondary structures in two conditions, and generated SP reactivities based on the simulated secondary structures in each condition using three types of empirical models representing different SP platforms, including two types of SHAPE reactivity^25, 35, 36^ and one type of icSHAPE reactivity^17^ (see Supplementary Methods). Biologically, it is often the case that a particular RNA molecule will occur as a mixture of structural conformations in a given cellular condition^37, 38^, so SVRs detected between conditions represent altered proportions comprising these mixtures of structural conformations. Thus, the between-condition differences in RNA secondary structures can be very subtle in reality. To mimic the complexity of the landscape of RNA secondary structures, we sampled multiple structural conformations from an ensemble of energy-function-based predictions, and mixed them in varying proportions in the simulated datasets (see Methods). The reactivities were thereafter simulated based on the pairing status of each nucleotide.

In detail, we sampled 10 conformations for each of the 100 RNA transcripts we randomly selected from transcripts in human embryonic kidney (HEK293) cells^17^. The lengths of the RNA transcripts ranged between 61 nt and 4,810 nt, and the simulated SVRs covered 3% to 55% of nucleotide positions of the RNA transcripts, with lengths between 1 nt and 16 nt. These data included a total of 13,162 simulated SVRs, with 6,587 being single nucleotide structural variations, 6,081 having lengths between 1 nt and 5 nt, and 494 SVRs with lengths greater than 5 nt (see Supplementary Fig. 4 for further details). Note that these simulated data echo the real world knowledge that the lengths of SVRs vary extensively^13, 19, 23, 24, 39^. We varied the strength of differential signals of simulated SVRs between “high”, “medium”, and “low”. Combining different levels of signal strength and the three types of generative models of reactivity, we have in total 9 simulation settings.

We compared the performance of DiffScan with two other SVR detection methods, deltaSHAPE and dStruct. Note that DiffScan adaptively determines the lengths of SVRs from data, whereas search length is a prespecified parameter for deltaSHAPE and dStruct. We set the search length to 5 nt for deltaSHAPE (the default setting suggested by the authors^22^) and to three different values (1 nt, 5 nt, and 11 nt) for dStruct.

To evaluate the accuracy of SVR boundaries mapping, we calculated the distance between each nucleotide position in the predicted SVRs and the true SVRs. In the ideal case, if a predicted SVR sits within a simulated SVR, the distance for each nucleotide position in the predicted SVR is zero. Oppositely, if a predicted SVR is off-target, *i*.*e*., containing many nucleotide positions that are neither in or close to the simulated SVRs, the distances for such nucleotide positions in the predicted SVR will be large.

The top-ranked SVRs identified by DiffScan are closer to the simulated SVRs than those by deltaSHAPE or dStruct in all 9 settings (Fig. 2a), demonstrating the superior performance of DiffScan in accurate identification of SVRs. Given that dStruct only considers nonoverlapping regions and relies on a pre-specified search length, and deltaSHAPE considers regions of fixed length, their performance is sensitive to whether or not pre-specified value of search length fits the actual lengths of the SVRs in a dataset. In contrast, DiffScan determines boundaries of SVRs by optimizing its scan statistic, thereby effectively distinguishing SVRs amongst overlapping regions of different lengths, leading to a finer level of granularity relative to deltaSHAPE and dStruct. Moreover, the superiority of DiffScan is consistent across different reactivity models and signal levels.

**Fig. 2.**
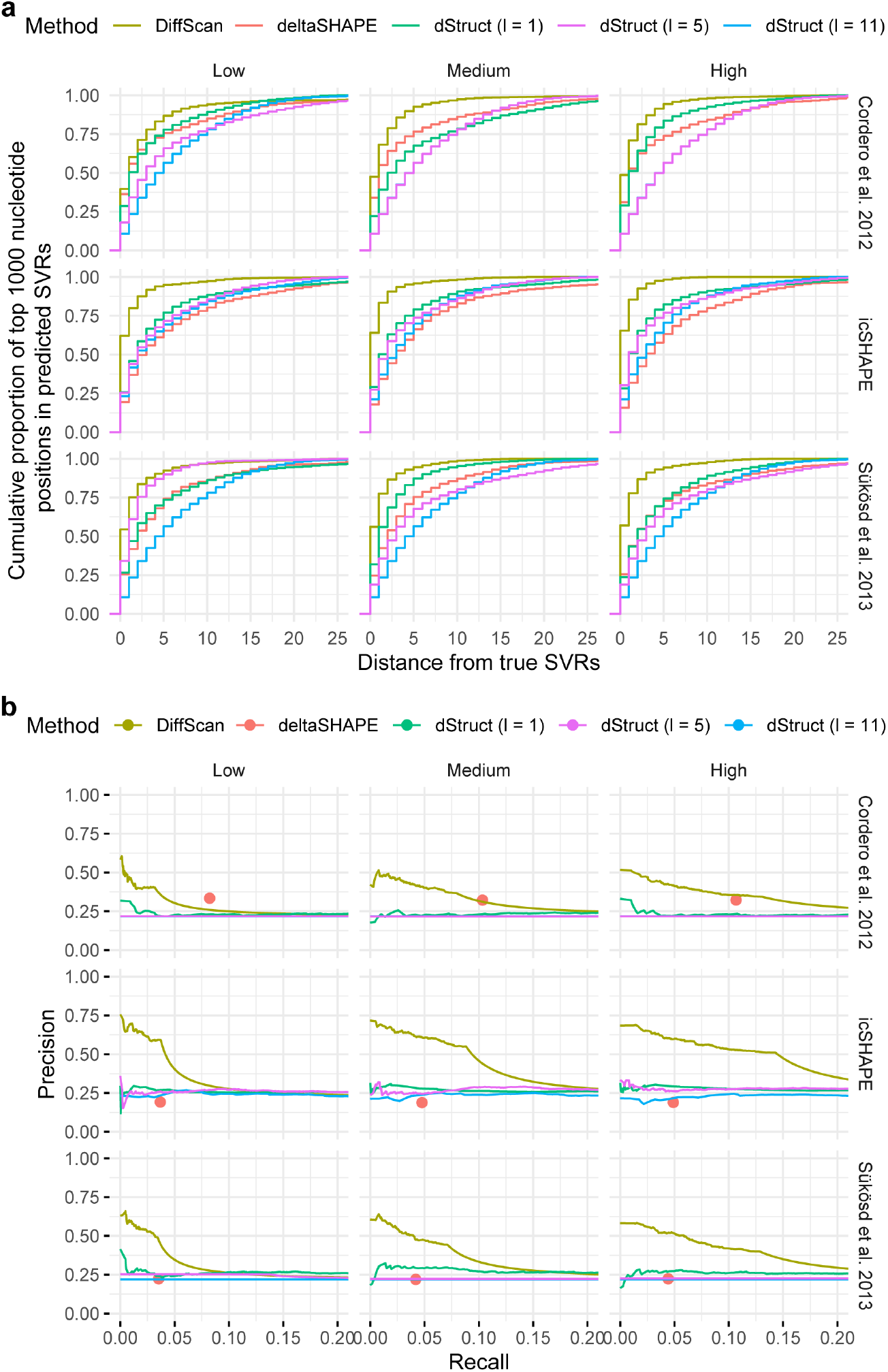
Comparison of DiffScan and other SVR detection methods with simulated datasets. For deltaSHAPE, we used the default search length of the method of 5 nt; for dStruct we used search length 1 nt, 5 nt, and 11 nt. **a** Distances between simulated SVRs and the top-ranked 1,000 nucleotide positions in predicted SVRs. **b** Precision-Recall curves for the prediction results from the various SVR detection methods. Rows: three types of reactivity generative models. Columns: three levels of strength of differential signals at simulated SVRs. Note deltaSHAPE does not allow external thresholding, and therefore it is represented as dots instead of curves.

We also evaluated the precision and recall rate at the nucleotide position level (see Methods) of the SVR detection methods. Among all three methods, DiffScan achieves the best precision at the same recall rate (Fig. 2b). Using optimization based on its scan statistic, DiffScan effectively identifies nonconsecutive SVR segments with nucleotide resolution. Although dStruct also sensitively identifies SVRs, it tends to output long, contiguous regions, which include a substantial proportion of nucleotide positions without structural variations, hampering the precision. The output of deltaSHAPE is a fixed set of regions that the threshold is internally decided, and it is represented as dots rather than curves in Fig. 2b. Although deltaSHAPE works well for the SHAPE reactivity model from Cordero *et al*.^35^, which is reasonable since the method was originally developed for SHAPE-MaP experiments, its precision is much lower than dStruct and DiffScan for other types of reactivity models.

### Sensitivity analysis

There are two tuning parameters in DiffScan: the radius of local windows (denoted by r) for smoothed calculation of positional differential signal, and the parameter associated with the penalty term (denoted by γ) in the scan statistic. In the above analysis of the simulated datasets, we set r to 2 nt and γ to 0.5 as their default values. We also conducted a sensitivity analysis wherein we tested DiffScan with the simulated datasets using eight other settings of (*r, γ*). In each setting, we calculated the mean precision of the predicted SVRs at recall values below 0.05. The coefficient of variation for the mean precisions corresponding to different tuning parameters is below 0.15 for all of the examined scenarios in the simulations (Supplementary Table 1), showing the robustness of DiffScan regarding to the tuning parameters. Notably, DiffScan consistently outperformed dStruct and deltaSHAPE in these tested parameter settings (Supplementary Fig. 5).

### Model validation with negative control datasets

To assess the extent of false positive discoveries of DiffScan and other SVR detection methods, we constructed six negative control datasets (Control 1-6, see Methods) from multiple SP platforms, including SHAPE-Seq and icSHAPE. In these datasets, no SVRs are present between the compared conditions. Thus, any significant SVRs identified by the detection methods would represent false positives. These comparisons included the PARCEL and RASA methods for the cases where raw read counts are available (Control 1, 2, 5, and 6). As dStruct requires within-condition replicates, it is not applicable for Control 5 and 6.

The false positive rates for DiffScan, dStruct, and PARCEL were below 0.05 for all tested datasets (Fig. 3a). It bears emphasis that the test space of DiffScan is tremendously larger than the other methods because it contains overlapping regions of differing lengths. Nevertheless, its false positive rates were well controlled. The error rate of RASA is slightly inflated in Control 2, but is well-controlled for the other datasets. Consistent inflation is observed for deltaSHAPE across different datasets, reporting false positive results covering ∼15% of nucleotide positions in the datasets. The fact that neither deltaSHAPE nor RASA control for the multiple testing problem can likely explain the observed inflation.

**Fig. 3.**
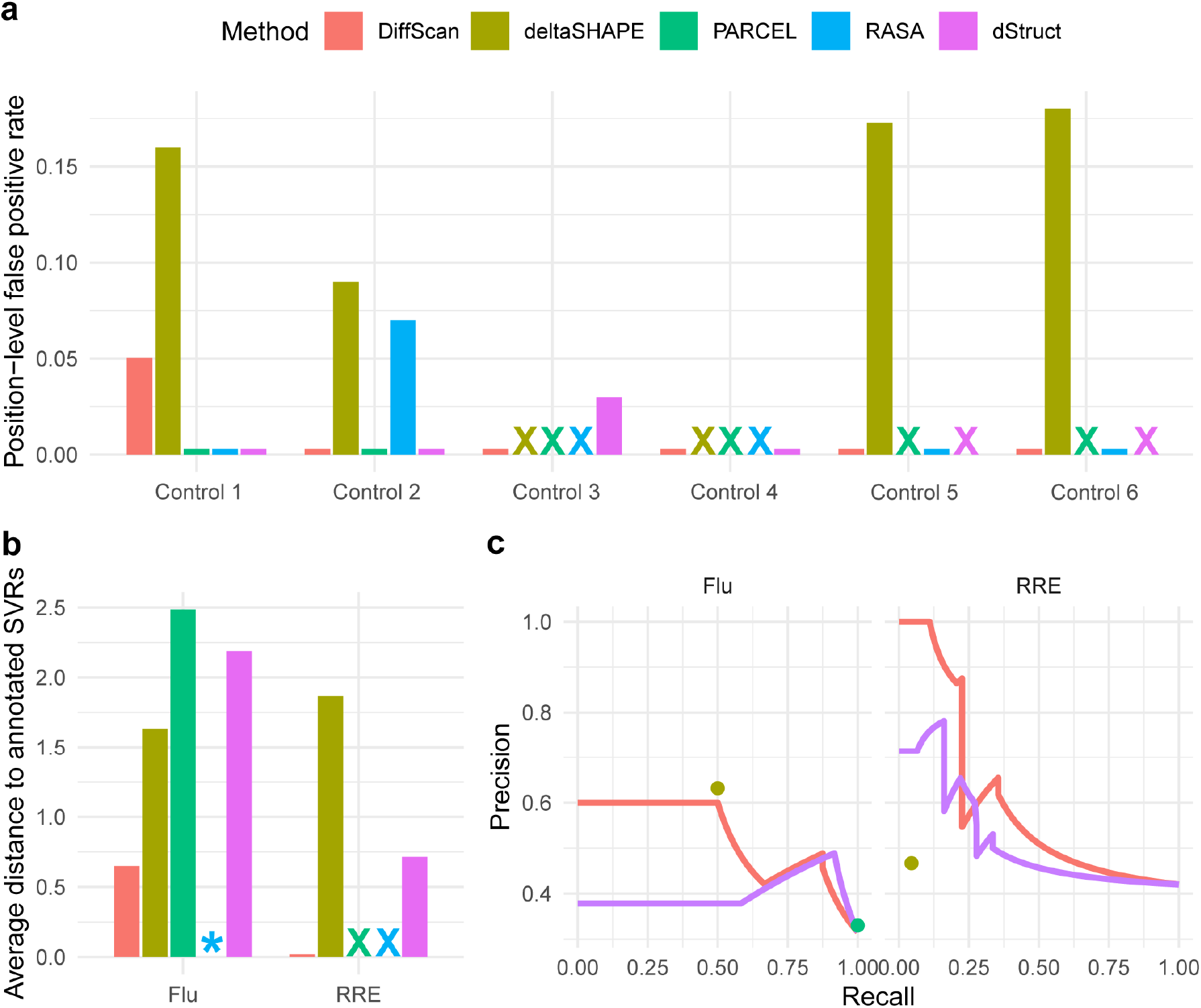
Comparison of DiffScan and existing SVR detection methods with negative control datasets and benchmark datasets. For deltaSHAPE, we used the default search length of the method of 5 nt; for dStruct we used search length 5 nt following the original article of the method. **a** Position-level false positive rate at a significance level of 0.05 in six negative control datasets (*i*.*e*., datasets having no SVRs). **b** Average distance from the top-10 ranked nucleotide positions in the predicted SVRs to the annotated SVRs in the benchmark datasets. **c** Precision-Recall curves for the benchmark datasets. deltaSHAPE and PARCEL are represented as dots. “X” indicates that the corresponding method was not applicable for the dataset. “*” indicates that the corresponding method did not report any region.

### Evaluation using benchmark datasets

We used two benchmark datasets with explicitly annotated SVRs between the compared conditions, curated by Choudhary *et al*.^23^ to support evaluation of different SVR detection methods. The Flu dataset^24^ is from a SHAPE-Seq experiment that measured the secondary structure of a *Bacillus cereus crcB* fluoride riboswitch *in vitro* in the presence or absence of fluoride. The transcript (100 nt in length) has five annotated SVRs, ranging in length between 1 nt and 8 nt. The RRE dataset^39^ is from a SHAPE-Seq experiment that measured the secondary structure of the HIV Rev-response element in the presence or absence of the Rev protein. There are seven annotated SVRs for Rev-RRE interactions in the transcript (369 nt in length), with lengths ranging between 7 nt and 39 nt.

We evaluated the average distance between predicted SVRs and the annotated SVRs and Precision-Recall curves output by DiffScan, deltaSHAPE, PARCEL, RASA, and dStruct (Fig. 3b-c). Consistent with our results when evaluating using simulated datasets and showcasing DiffScan’s superior performance for accurately locating SVRs, DiffScan achieved the lowest average distance from the annotated SVRs of the benchmark datasets. For the Flu dataset, DiffScan identified all the five annotated SVRs with their boundaries accurately mapped (Supplementary Fig. 6). In contrast, PARCEL predicted a long region, covering 72% of all nucleotide positions in the transcript, that spanned all five annotated SVRs. However, it did not distinguish annotated SVRs from flanking nucleotides between SVRs and falsely entailed many nucleotide positions (67% of the nucleotide positions in the predicted SVRs) without structural variations. Similarly, dStruct identified a long region covering 37% of all nucleotide positions, which covered two annotated SVRs, while 62% of the nucleotide positions in the predicted SVRs had no structural variation. RASA did not report any significant region for the Flu dataset. deltaSHAPE reported six regions, missing an annotated SVR while containing a false positive discovery. For the RRE dataset, DiffScan detected three regions which are located within three annotated SVRs. PARCEL and RASA were not applicable for the RRE dataset because raw count data were not available. deltaSHAPE and dStruct both reported four regions, which overlapped with two and four annotated SVRs respectively. Compared to DiffScan, they reported regions with much larger average distance to annotated SVRs, since they included many nucleotide positions (53% and 35% of the nucleotide positions in the predicted SVRs for deltaSHAPE and dStruct) without structural variation.

In terms of precision and recall measures, DiffScan also achieved the best area under the curve (AUC) values with both datasets (Fig. 3c). Note that because deltaSHAPE and PARCEL do not allow threshold tunings, so they are represented as dots rather than curves in Fig. 3c. deltaSHAPE performed comparably to DiffScan for the Flu dataset. However, the precision of deltaSHAPE was much lower than DiffScan and dStruct for the RRE dataset, since the predicted SVRs of deltaSHAPE included many nucleotide positions without structural variation. In general, dStruct had lower precision compared to DiffScan when the recall rate was set to the same level, consistent to the trends we observed when analyzing the simulated datasets.

### Roles of SVRs in regulating mRNA transcripts for mitochondria-associated proteins

We applied DiffScan to a recently reported icSHAPE dataset that mapped RNA secondary structure across human cellular compartments covering chromatin (Ch), nucleoplasm (Np), and cytoplasm (Cy)^17^. The DiffScan-predicted SVRs for the Ch versus Np (Fig. 4a) and for the Np versus Cy (Supplementary Fig. 7) comparisons mostly involved protein binding sites and RNA modification sites. As we also had data for RNA abundance in the Ch, Np, and Cy samples, we were interested in the potential impacts of SVRs on regulating mRNA abundance. Comparison of mRNA abundance in the Np versus Cy samples clearly placed the mRNAs into two subgroups, with mRNA levels in subgroup II decreased sharply in the Cy fraction (Fig. 4b). A manual check revealed that all 12 mRNAs of subgroup II encode mitochondria-associated proteins. In addition, DiffScan identified SVRs in all of these 12 transcripts (p value = 8.4e-3), suggesting an association between RNA structural variation and the abundance of mRNAs encoding mitochondria-associated proteins.

**Fig. 4.**
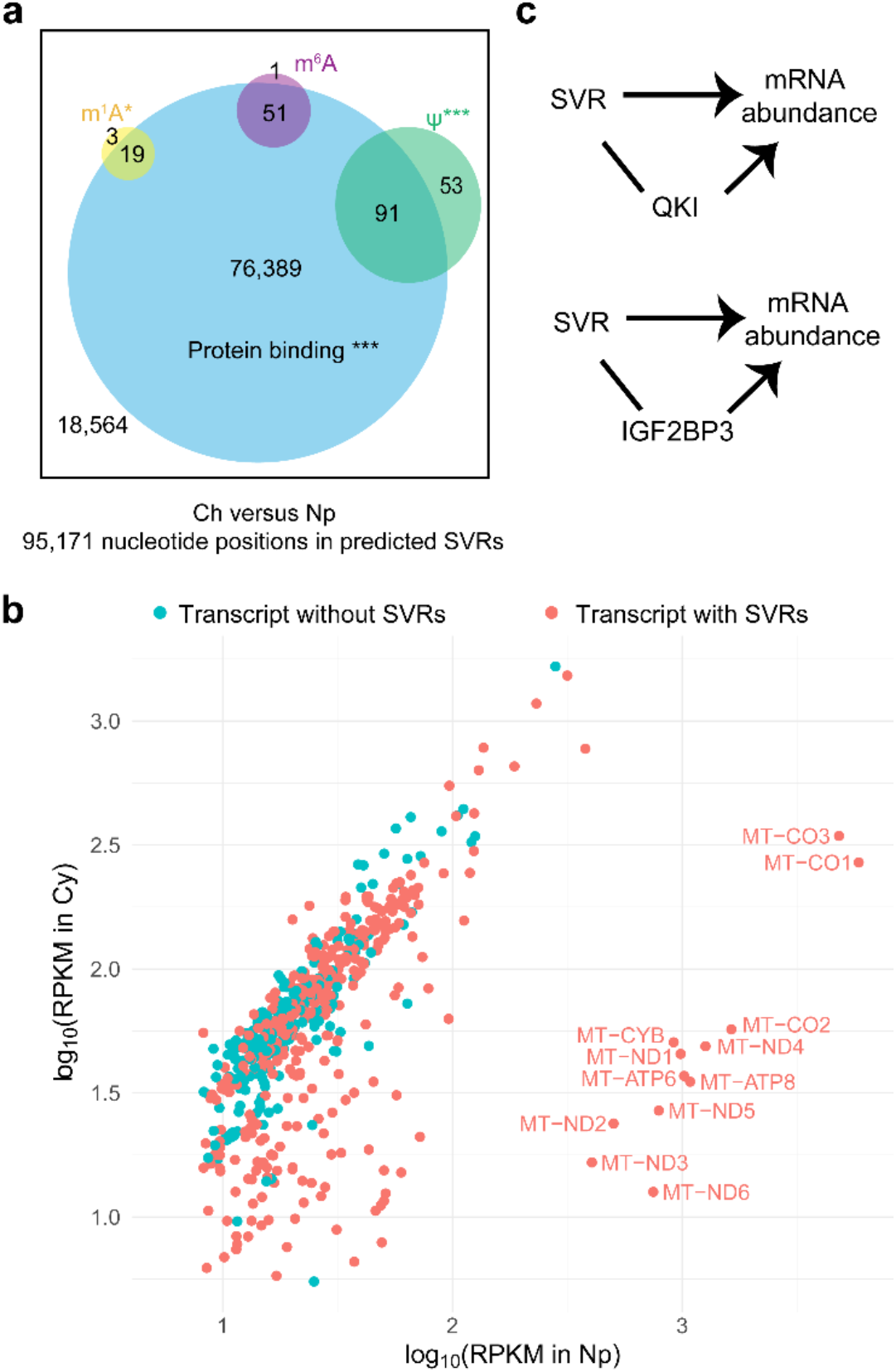
Application of DiffScan to explore the roles of RNA structural variation in regulation of mRNA abundance in an icSHAPE dataset mapping RNA structure across human cellular compartments. Ch: chromatin, Np: nucleoplasm, Cy: cytoplasm. **a** Predicted SVRs between Ch and Np were enriched with protein binding sites and RNA modification sites. *p value (Fisher’s exact test) < 0.05, ***p value < 1e-6. **B** RPKM of transcripts and the prediction results of DiffScan for Np versus Cy. **c** Statistical inference supports the direct roles of SVRs in regulation of mRNA abundance.

To further investigate this association, we used the FIMO module from the MEME suite^40^ to search for RBPs with binding motifs enriched in the predicted SVRs from the Np versus Cy comparison (see Methods). Three enriched RBPs were identified, including quaking (QKI), insulin-like growth factor 2 mRNA-binding protein 3 (IGF2BP3), and polyadenylate-binding protein 1 (PABPC1). QKI had two significantly enriched motifs, ACUAACA (corrected p value = 1.4e-5) and UACUAAC (corrected p value = 7.8e-4). QKI predominantly localizes in the Np^41, 42^, and it has been reported to regulate various biological processes such as myelinization^43^, mRNA stability, and mRNA export^42^. Of particular note, a recent study reported that QKI binds mRNAs encoding mitochondrial proteins and transcriptional coactivators known to regulate mitochondrial biogenesis and function: QKI decreases the stability and nuclear export of these RNAs^44^. Therefore, the binding of QKI to the mRNA transcripts of subgroup II, which all encode mitochondria-associated proteins, may in part explain their reduced abundance in the Cy samples.

Considering the biological evidence and the potential roles of SVRs, we enumerated four possible causal models to describe the relationships among SVRs, QKI binding, and changes in mRNA abundance (Model 1-4 in Supplementary Fig. 8; see Methods). In Model 1, QKI binding causes SVRs in the mRNA transcripts of subgroup II and also regulates their abundance in a separate pathway, and thus SVRs have no contribution to the observed decline of mRNA abundance in the Cy. In Model 2-4, SVRs either directly or indirectly (mediated by QKI binding) regulate mRNA abundance. The observed data rejects Model 1 and Model 2 (p value = 2.5e-91), and therefore supports the direct roles of SVRs in regulation of mRNA abundance. However, Model 3 and 4 are statistically equivalent, *i*.*e*., not distinguishable. In another word, SVRs have direct effect in regulating mRNA abundance, in addition to QKI binding (Fig. 4c); however, the causal relationships between SVR and QKI binding cannot be determined with current data.

The second RBP, IGF2BP3, had one motif (ACAAACA) enriched among the predicted SVRs (corrected p value = 5.4e-3). IGF2BP3 is almost exclusively cytoplasmic, and has been conceptualized to “cage” its target mRNAs into cytoplasmic protein-RNA complexes as they enter the Cy^45^. Recently, the upregulation of IGF2BP3 has been associated with mitochondrial dysfunction^46^. We reason that IGF2BP3 cages mRNA transcripts encoding mitochondria-associated proteins as they enter the Cy, accounting for the observed decline in the abundance of such mRNA transcripts. We enumerated possible causal models (Supplementary Fig. 8) and performed hypothesis testing to explore the relationships among SVRs, IGF2BP3 binding, and changes in mRNA abundance. According to our inference, there is a direct effect from the detected SVRs to the changes in mRNA abundance (Fig. 4c; p value = 7.4e-49), although it is not clear whether the SVRs’ regulation of mRNA abundance is also mediated by IGF2BP3 binding.

In summary, the DiffScan-predicted SVRs provide insight into the functional roles of RNA secondary structure and structural variation in regulating mRNAs encoding mitochondria-associated proteins.

## Discussion

We have developed DiffScan, a computational framework for differential analysis of RNA secondary structure measured in multiple SP platforms. Our method provides several advantages. First, it adaptively estimates the lengths and locations of SVRs with single-nucleotide resolution. Compared to existing SVR detection methods, predicted SVRs by DiffScan are closer to true SVRs, both from simulated and benchmark datasets. Second, DiffScan is compatible with multiple SP platforms with robust performance. Existing SVR detection methods are usually tailored for a specific SP platform, and their generalization to other SP platforms can be prohibitive. Our method flexibly accommodates multiple SP platforms through its Normalization module and using the nonparametric test we implemented. This enables flexible analysis that is adaptable to suitable data types reflecting particular biological problems of interest. For example, the SHAPE-Seq platform with fast-acting reagent is suitable for *in vitro* studies of RNA folding dynamics^47^, while the icSHAPE platform utilizing slow-acting reagent allows *in vivo* transcriptome-wide structure probing^16, 17^. As demonstrated with simulated and benchmark datasets, the superior performance of DiffScan in terms of statistical power and accuracy for identifying SVRs is robust across different SP platforms. Third, DiffScan rigorously controls for the family-wise error rate in SVR detection, which is particularly influential for transcriptome level analyses. At last, the SVR simulation framework we devised aptly reflects the complexity of SVR detection problems in the real world. We have developed a “simulation” module in our software, which can facilitate evaluation of more upcoming SVR detection methods in the future.

DiffScan has several limitations. First, the Normalization module in DiffScan relies on a structurally invariant set of nucleotide positions, which is identified via a built-in data-driven strategy. The strategy would fail in extreme cases when almost all nucleotide positions in the studied transcripts exhibit structural variations, although we expect it is rarely the case in practice^4, 16, 17, 48^. In addition, when the structurally invariant set can be specified by prior knowledge, the normalization step can be easily adapted. Second, we recognize the statistical power of DiffScan is not optimized for a particular SP platform, so when the distribution of reactivities is known *a priori*, use of an appropriate parametric test would yield increased power. Third, we note that the ultimate goal of studying SVRs is to understand the functional roles of RNA structure, which often involves cross-examination of other data sources, including motif analysis of RBPs, multi-omics datasets, *etc*. Incorporation of these multi-source data may help to accurately annotate SVRs^11^.

## Acknowledgements

This work was supported by the National Natural Science Foundation of China (Grants No. 91740204 and 91940306) and the State Key Research Development Program of China (Grant No. 2019YFA0110002) to Q.C.Z. LH acknowledges research support from the National Science Foundation of China (Grant No. 12071243).

## Author contributions

B.Y., Q.C.Z., and L.H. conceived the study and wrote the manuscript. B.Y. and L.H. developed the computational framework. B.Y. and P.L. performed the data preparation and statistical analysis. B.Y., Q.C.Z., and L.H. interpreted the results. All authors approved the manuscript.

## Methods

### Structure Probing Datasets

We evaluated the power of DiffScan and other SVR detection methods in two SHAPE-seq datasets, of which the SVRs were curated in Choudhary *et al*.^23^. The Flu dataset probed the *Bacillus cereus crcB* fluoride riboswitch *in vitro* with and without fluoride ions, with four replicates for each condition. The transcript is 100 nt in length, and nucleotide positions 12-17, 22-27, 38-40, 48 and 67-74 were identified as SVRs between conditions. The RRE dataset studied the HIV Rev-response element, with three replicates measured in the presence and absence of Rev. The Rev-RRE interaction sites are considered SVRs. We obtained raw read counts and reactivities for the Flu dataset and reactivities for the RRE dataset, based on data availability.

Synthetic negative control datasets were constructed from real SP data to assess the specificity of DiffScan and existing methods. For construction, we randomly split replicates in a single condition into two groups, and identified SVRs by contrasting the two groups. Any reported SVRs were considered false positive results. In particular, control datasets 1-4 were obtained by randomly splitting samples in the Flu dataset in the absence (Control 1) and presence (Control 2) of fluoride, and the RRE dataset with (Control 3) and without Rev (Control 4). To include more SP platforms in evaluation, we constructed another two control datasets from an icSHAPE dataset of mouse SRP RNA (272 nt)^10^, by contrasting the two *in vitro* samples (Control 5) and the two *in vivo* samples (Control 6).

To apply DiffScan in transcriptome level analysis, we downloaded the icSHAPE dataset^17^ in Sun *et al*., which mapped RNA secondary structure across HEK293 cellular compartments including chromatin, nucleoplasm, and cytoplasm. Transcripts with RPKM less than 5 or RT stop less than 2 for all nucleotide positions were excluded^17, 49^. Then nucleotide positions with coverage less than 200 were removed. As a result, 1,352 transcripts (covering 826,917 nucleotide positions) were compared between chromatin and nucleoplasm, and 1,851 transcripts (covering 1,220,273 nucleotide positions) were compared between nucleoplasm and cytoplasm.

### Normalization module

Suppose that there are n_A_ replicates from condition A, and the reactivity of nucleotide position j in replicate i is 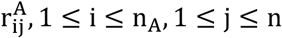, where n is the length of the transcript. Similarly, for n_B_ replicates from condition B, the reactivity of nucleotide position j in replicate k is 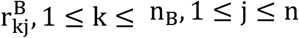. We normalized reactivities using the following steps.

Step 1: 90% winsorization is applied to the reactivities in each replicate separately to remove outliers, *i*.*e*., the bottom 5% of reactivities are set to the 5th percentile while the upper 5% of reactivities are set to the 95th percentile. After that, reactivities are scaled into range [0,1] by subtracting the minimum and then dividing the result by the new maximum.

Step 2: Quantile normalization is applied to within-condition replicates to match the quantiles, which is frequently used in normalization analysis of genome-wide assays^30, 50^.

Step 3: To make between-condition reactivities comparable, we first determine a structurally invariant set *S, i*.*e*., nucleotide positions that do not exhibit structural variation across conditions. The reactivities are normalized so that the adjusted reactivities are similar between conditions in the structurally invariant set. The determination of *S* is described in Supplementary Methods. Then we learn a linear transformation rule from *S*, which converts reactivities of one condition to the same level of those of the other condition in *S*, and extrapolate the transformation to all nucleotide positions to obtain normalized reactivities.

We learn how to transform between-condition reactivities to the same level in *S* by fitting the following robust regression utilizing iterated re-weighted least squares with Huber’s *M* estimate^51^. (For the sake of simplicity, we will still use 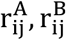 to denote reactivities processed by Step 1 and Step 2.)

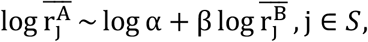

where log α corrects for the difference in sequencing depth and β corrects for the difference in signal-to-noise ratio. Given that within-condition replicates are normalized by quantile normalization, it follows that 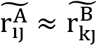 in *S* if we transform 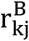 into 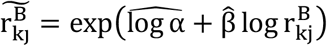 and let 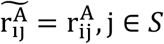, j ∈ *S*. Based on this, we extrapolate the fitted model to all nucleotide positions to obtain normalized reactivities 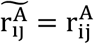 and 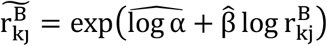 1 ≤ j ≤ n.

### Differential Analysis at the Position Level

With normalized reactivities as input, we first test position-level differences between conditions to derive positional p values. To encompass various SP platforms, a nonparametric Wilcoxon test is used, which evaluates the structural variation at nucleotide position j by contrasting reactivities between conditions in a small window surrounding j. Technically, we define

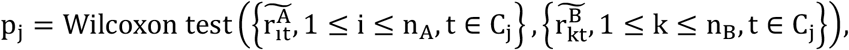

where 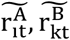 are normalized reactivities, C_j_ is a small window centering at nucleotide position j with radius r, and p_j_ is the two-sided p value. By default, we set r to 2 nt.

### Summarizing Position-level Signals via Scan Statistics

To concatenate position-level signals into regional signals, a scan statistic is defined for each region R,

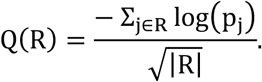

With penalty of region length |R|, extending a candidate region will accumulate position-level signals at the expense of penalization. As a result, SVRs with accurately mapped boundaries can be distinguished based on the magnitude of the scan statistic.

### Statistical Inference for Detection of SVRs

In view of the dynamic locations and lengths of SVRs, we scan the transcripts by enumerating all candidate regions with suitable lengths, *i*.*e*., every region in ℛ = {R | L_min_ ≤ |R| ≤ L_max_}. To determine whether a region R is selected as an SVR, we test

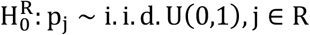

based on the magnitude of Q(R). We implemented a Monte Carlo approach to control the family-wise error rate for DiffScan (see Supplementary Methods).

### Simulations

We simulated reactivities for two conditions through the following three steps.

Step 1. 100 human RNA sequences in an icSHAPE dataset^17^ were randomly selected. For each RNA sequence, we sampled 10 conformations from the Boltzmann distribution of secondary structure^52^ utilizing the RNAsubopt program in ViennaRNA version 2.4.15^53^.

Step 2. Conditional on the pairing status of each nucleotide in a transcript, the reactivities can be sampled from pre-trained reactivity distributions. Three distributions were used: Cordero *et al*.^35^ and Sükösd *et al*.^36^ as fitted from SHAPE data, and also reactivity distributions we fitted from icSHAPE data (Supplementary Methods).

Step 3. Different biological conditions were characterized by differential compositions of the 10 conformations generated in Step 1. The reactivities of each conformation were linearly combined accordingly to generate the observed reactivities in each condition. Specifically, for condition A, the 10 conformations of a transcript were assigned with weights 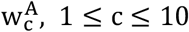, with 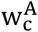 corresponding to the weight of the c^th^ conformation in condition A, with 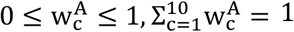. For condition B, the weight of the c^th^ conformation was 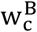, with 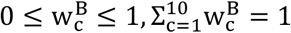. To simulate SVRs that cover a reasonable proportion (approximately 20%^16, 17^) of all nucleotide positions of the 100 transcripts, we allocated 90% of the weight to the first two conformations and randomly distributed the remaining 10% to the other eight conformations. We then generated reactivities in condition A and B separately by changing the weights of the first two conformations. Thus, nucleotide positions with different pairing status between the first two conformations form SVRs. Furthermore, replicates were simulated by adding random noise (N(0,0.1^2^)) to the weights of the first two conformations. Finally, simulations were conducted at three levels of signal strength respectively by setting 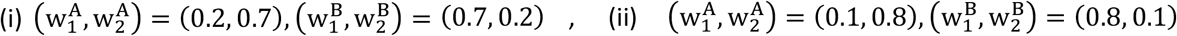, and 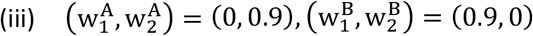.

### Performance Metrics for Validation of DiffScan

In the simulated datasets, we evaluated the performance of different methods with two metrics. First, to evaluate how accurately simulated SVRs (*i*.*e*., true SVRs to detect between the compared conditions in the simulated datasets) were mapped by predicted SVRs by a method, we took out 1,000 nucleotide positions in the top-ranked SVRs by the method. The distance from each of the nucleotide positions to simulated SVRs was calculated. Second, we calculated the Precision-Recall curve. At varying significance levels, the value of precision was calculated as the number of correctly predicted nucleotide positions divided by the number of all predicted nucleotide positions. The value of recall rate was calculated as the number of correctly predicted nucleotide positions divided by the number of all nucleotide positions in simulated SVRs.

In the negative control datasets (Control 1-6), to access whether the false positive discoveries of a method can be controlled at a nominal level, we calculated the position-level false positive rate as the number of predicted nucleotide positions at significance level 0.05 divided by the length of the transcript.

In the benchmark datasets with annotated SVRs (Flu and RRE)—similar to our processing of the simulated datasets—we calculated (1) the average distance from 10 nucleotide positions in top predicted SVRs to annotated SVRs and (2) the Precision-Recall curve.

### Implementation of existing methods

The software deltaSHAPE version 1.0 available at the Weeks lab website^22^ was utilized to implement deltaSHAPE. As far as we know, there is no official software release for PARCEL or RASA; and we implemented them utilizing custom scripts from existing literature^23^. The R package dStruct version 1.0.0 was utilized to implement dStruct.

### Application to icSHAPE data

#### Reactivity calculation

To calculate reactivities, we assumed that RT_j_ ∼ Binomial(N_j_, p_j_), in which RT_j_ is the RT stop at position j in the case experiment, N_j_ is the number of times that position j is exposed to the probing molecules, and p_j_ is the probability that position j is modified. The maximum likelihood estimator for p_j_ is 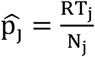. On the other hand, the coverage at position j in the control experiment 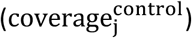 is approximately proportional to N_j_. Therefore, we calculated the reactivity at position j as 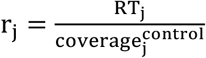^54^. Based on this, two count replicates from case experiments were paired with two count replicates from control experiments to produce two reactivity replicates under each condition.

#### Enrichment analysis of RBP binding motifs

We collected 154 RBP binding motifs from the CISBP-RNA Database^55^ and the ATtRACT database^56^. For each motif, we searched in the predicted SVRs for significant motif hits utilizing the FIMO module (--norc --thresh 0.001) from the MEME suite^40^, and the number of significant hits in each SVR were counted.

To evaluate the significance of motif enrichment, we randomly sampled null regions in non-SVR regions in the same transcripts, with region length matched to the predicted SVRs. The number of significant hits in each null region is recorded. After that, we performed a one-sided Wilcoxon signed-rank test for the two count vectors of significant hits to evaluate the significance of enrichment. Motifs with corrected p value (Benjamini-Hochberg method) less than 0.05 were considered significantly enriched. We further consolidated the result by filtering out motifs that lost enrichment, *i*.*e*., with corrected p value (Benjamini-Hochberg method) greater than 0.01, when the null regions were sampled from the non-SVR regions of all transcripts with SVRs.

#### Inference for the relationships among SVRs, RBP binding, and mRNA abundance

We considered four models of which each explicitly describes the relationships among SVRs, changes in RBP binding, and changes in mRNA abundance (Model 1-4 in Supplementary Fig. 8). For model selection, three variables of each mRNA transcript in the Np versus Cy comparison were used: (1) whether DiffScan predicted any SVRs in the Np versus Cy comparison for the transcript (*svr; svr* = “yes” or “no”), which quantifies its structural variations between Np and Cy, (2) log_10_(fold change of RPKM; Cy/Np) of the transcript (*log10*-*fold-change-of-RPKM*), which quantifies the change in its abundance between Np and Cy, and (3) a *change-in-RBP-binding* variable quantifying the changes in the binding of a specified RBP between Np and Cy for the transcript. Because QKI predominantly localizes in the Np, the *change-in-RBP-binding* variable for QKI of a transcript can be approximated by a score quantifying the level of QKI binding in the Np for the transcript. To calculate the score, we searched in the transcript for significant hits of QKI motifs utilizing the FIMO module (--norc --thresh 0.001), and the number of significant motif hits was taken as the score quantifying the level of QKI binding in the Np for the transcript. Similar processing was implemented for IGF2BP3, because it is almost exclusively cytoplasmic.

Given the three variables of each transcript, *i*.*e*., *svr, log10-fold-change-of-RPKM*, and *change-in-RBP binding* for a specified RBP, we tested conditional independence between *svr* and *log10-fold-change-of-RPKM* given *change-in-RBP-binding* utilizing the conditional independence test for hybrid Bayesian networks implemented in the bnlearn R package^57^. In Model 1-2, *svr* and *log10-fold-change-of-RPKM* are conditionally independent given *change-in-RBP-binding*, whereas *svr* regulates *log10-fold-change-of-RPKM* given *change-in-RBP-binding* in Model 3-4. In cases where Model 1-2 are rejected by the conditional independence test, Model 3-4 are supported. However, Model 3 and Model 4 are statistically equivalent with current data.

## Data availability

All data used in the manuscript were publicly available. The benchmark datasets were downloaded from https://doi.org/10.5281/zenodo.2536501. The icSHAPE dataset for transcriptome analysis were downloaded from the GEO database with accession code GSE117840 (https://www.ncbi.nlm.nih.gov/geo/query/acc.cgi?acc=GSE117840).

## Code availability

All codes and the developed software were publicly available at https://github.com/yub18/DiffScan.

