## Supplementary Information for "Differential Analysis of RNA Structure Probing Experiments at Nucleotide Resolution: Uncovering Regulatory Functions of RNA Structure"

### Supplementary Note

#### Normalization module: Determination of the structurally invariant set *S*

*S* is determined as the nucleotide positions with reactivities uniformly high or uniformly low across all replicates. These positions are probably simultaneously paired or simultaneously unpaired in both conditions. Technically, given two parameters $(\mathrm{l},\mathrm{u}), 0 \leq \mathrm{l} \leq 0.5 \leq \mathrm{u} \leq 1$,

$$S_i^{A,high} = \left\{ j \middle| r_{ij}^{A} \geq \mathrm{quantile}\left(\left\{r_{it}^{A}, 1 \leq t \leq n\right\};\ u\right)\right\}$$

consists of nucleotide positions with high reactivities in replicate $\mathrm{i}$ from condition A, and

$$S_k^{B,high} = \left\{ j \middle| r_{kj}^{B} \geq \mathrm{quantile}\left(\left\{r_{kt}^{B}, 1 \leq t \leq n\right\};\ u\right)\right\}$$

consists of nucleotide positions with high reactivities in replicate $\mathrm{k}$ from condition B. Therefore,

$$S^{high} = \left(\cap_{i=1}^{n_A} S_i^{A,high}\right) \cap \left(\cap_{k=1}^{n_B} S_k^{B,high}\right)$$

consists of nucleotide positions with uniformly high reactivities across all replicates. Similarly,

$$S^{low} = \left(\cap_{i=1}^{n_A} S_i^{A,low}\right) \cap \left(\cap_{k=1}^{n_B} S_k^{B,low}\right)$$

consists of nucleotide positions with uniformly low reactivities across all replicates, where

$$S_i^{A,low} = \left\{ j \middle| r_{ij}^{A} \leq \mathrm{quantile}\left(\left\{r_{it}^{A}, 1 \leq t \leq n\right\};\ l\right)\right\}$$

and

$$S_k^{B,low} = \left\{ j \middle| r_{kj}^{B} \leq \mathrm{quantile}\left(\left\{r_{kt}^{B}, 1 \leq t \leq n\right\};\ l\right)\right\}.$$

Then the invariant set is determined as

$$S = S^{high} \cup S^{low}.$$

At last, $(\mathrm{l},\mathrm{u})$ is selected by solving the following optimization problem with grid searching.

$$(\mathrm{l},\mathrm{u}) = \underset{\substack{0.05 \leq l \leq 0.4 \\ 0.6 \leq u \leq 0.95 \\ |S| \geq \max\left(10, \frac{n}{20}\right)}}{\mathrm{argmax}} \ \text{spearman correlation}\left(\overline{r_j^A}, \overline{r_j^B}\right)_{j \in S}.$$

The idea is that when nucleotide positions with structural variations are added to *S*, the spearman correlation of between-group reactivities in *S* should decrease.

### Scan module: Monte Carlo approach controlling for family-wise error rate

At significance level $\alpha$, we first calculate a threshold $h_\alpha$ for the scan statistic $Q(R)$ from Monte Carlo sampling. Then we implement the following algorithm which outputs the predicted SVRs by DiffScan.

```
Input: 𝓡 = {L_min ≤ |R| ≤ L_max}, {Q(R′) | R′ ∈ 𝓡}.
1:  SVR ← ∅.
2:  while 𝓡 ≠ ∅ do
3:     R ← argmax_{R′∈𝓡} Q(R′).
4:     if Q(R) < h_α then
5:        return SVR
6:     else
7:        SVR ← SVR ∪ {R}.
8:        𝓡 ← {R′ ∈ 𝓡 | R′ does not overlap with R}.
9:     end if
10: end while
11: return SVR
```

To calculate the threshold $h_\alpha$, we sample $\widetilde{p_j} \sim \mathrm{i.i.d.}\, U(0,1), 1 \le j \le n$ and compute the corresponding scan statistic values

$$\widetilde{Q(R)} = \frac{-\sum_{j\in R}\log\left(\widetilde{p_j}\right)}{\sqrt{|R|}}, R \in \mathcal{R}$$

and the extreme statistic

$$\widetilde{Q_{max}} = \max\left\{\widetilde{Q(R)} \mid R \in \mathcal{R}\right\}.$$

This process is repeated N times to get N replications of $\widetilde{Q_{max}}$, from which we calculate

$$h_\alpha = \mathrm{quantile}\left\{\widetilde{Q_{max}}; 1-\alpha\right\}.$$

In practice, we cut the transcript and/or the transcriptome into 100 nt segments and then enumerate contiguous regions with a minimum length $L_{min} = 1$ nt and a maximum length $L_{max} = 20$ nt in the 100 nt segments. Note that $L_{max} = 20$ nt is not the upper bound of length of finally predicted SVRs, since DiffScan would predict multiple SVRs which might adjoin each other within long SVRs.

**Simulations: three types of reactivity models used to simulate reactivities**

Two types of distributions of SHAPE reactivities were fitted in an existing literature[1] from two independent sources.

Cordero et al.[2]:

$$\text{reactivity}_{\text{paired}} \sim \text{Generalized Extreme Value (GEV) distribution}(\mu = 0.0947, \sigma = 0.0672, \epsilon = 0.2352),$$

$$\text{reactivity}_{\text{unpaired}} \sim \text{GEV}(\mu = 0.2198, \sigma = 0.1852, \epsilon = 0.5426).$$

Sükösd et al.[3]:

$$\text{reactivity}_{\text{paired}} \sim \text{GEV}(\mu = 0.0523, \sigma = 0.0680, \epsilon = 0.8681),$$

$$\text{reactivity}_{\text{unpaired}} \sim \text{exponential}(\lambda = 1.4638).$$

We fitted reactivity distributions for paired and unpaired nucleotides characterizing the statistical nature of reactivities acquired using the icSHAPE platform utilizing reactivities of 100 transcripts we selected from an icSHAPE dataset[4]:

$$\text{reactivity}_{\text{paired}} \sim \pi_1 \delta_0 + (1 - \pi_1) \exp\left(\text{Normal}(\mu_1, \sigma_1^2)\right),$$

$$\text{reactivity}_{\text{unpaired}} \sim \pi_2 \delta_0 + (1 - \pi_2) \exp\left(\text{Normal}(\mu_2, \sigma_2^2)\right),$$

in which $\delta_0$ is a point mass at 0 and $\widehat{\pi_1} = 0.68, \widehat{\mu_1} = -3.40, \widehat{\sigma_1^2} = 3.53, \widehat{\pi_2} = 0, \widehat{\mu_2} = -2.69, \widehat{\sigma_2^2} = 0.97$. ($\mu_1, \sigma_1^2$ and $\mu_2, \sigma_2^2$ were estimated using a two-component Gaussian finite mixture model[5] from the nonzero values of the icSHAPE reactivities. $\pi_1$ and $\pi_2$ were further estimated combined with the frequency of paired and unpaired nucleotides in the simulated secondary structures.)

### Supplementary Tables

| Coefficient of variation | | Level of strength of differential signals in SVRs | | |
|---|---|---|---|---|
| | | Low | Medium | High |
| Type of reactivity distributions to simulate reactivities | Cordero et al. | 0.133 | 0.059 | 0.059 |
| | icSHAPE | 0.147 | 0.068 | 0.086 |
| | Sükösd et al. | 0.148 | 0.082 | 0.083 |

**Supplementary Table 1: Sensitivity analysis of DiffScan in terms of tuning parameters.** For each scenario in the simulations, the coefficient of variation for nine values of mean precision at recall values below 0.05, corresponding to the nine settings of $(r, \gamma)$: $r \in \{1, 2, 3\}$ and $\gamma \in \{0.4, 0.5, 0.6\}$, is displayed.

### Supplementary Figures

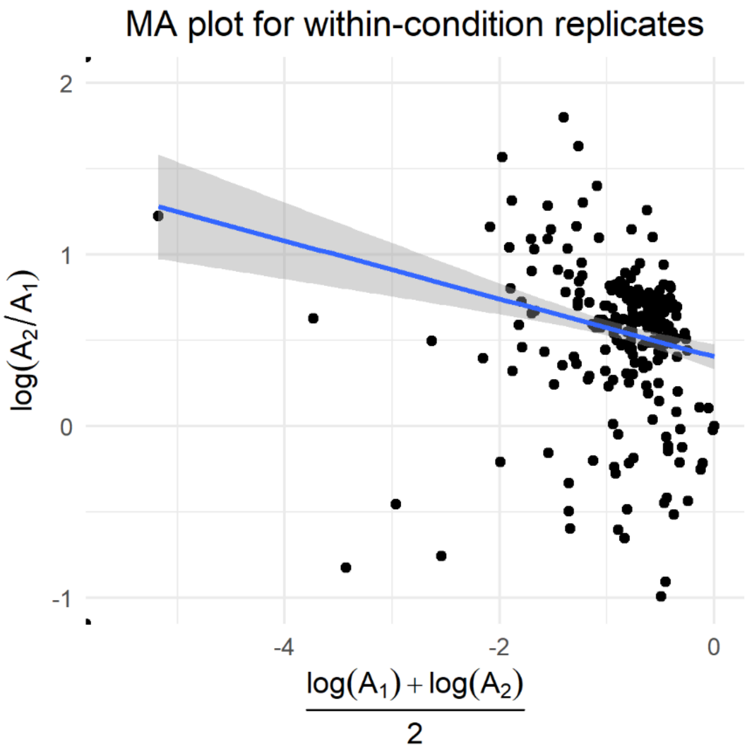


**Supplementary Figure 1: Systematic bias between reactivity replicates.** $\mathrm{A}_1, \mathrm{A}_2$ are within-condition replicates of the SRP *vivo* dataset (*i.e.*, Control 6; see Materials and Methods). The fitted line should approximate $\log\left(\frac{\mathrm{A}_2}{\mathrm{A}_1}\right) = 0$ if $\mathrm{A}_1, \mathrm{A}_2$ are comparable.

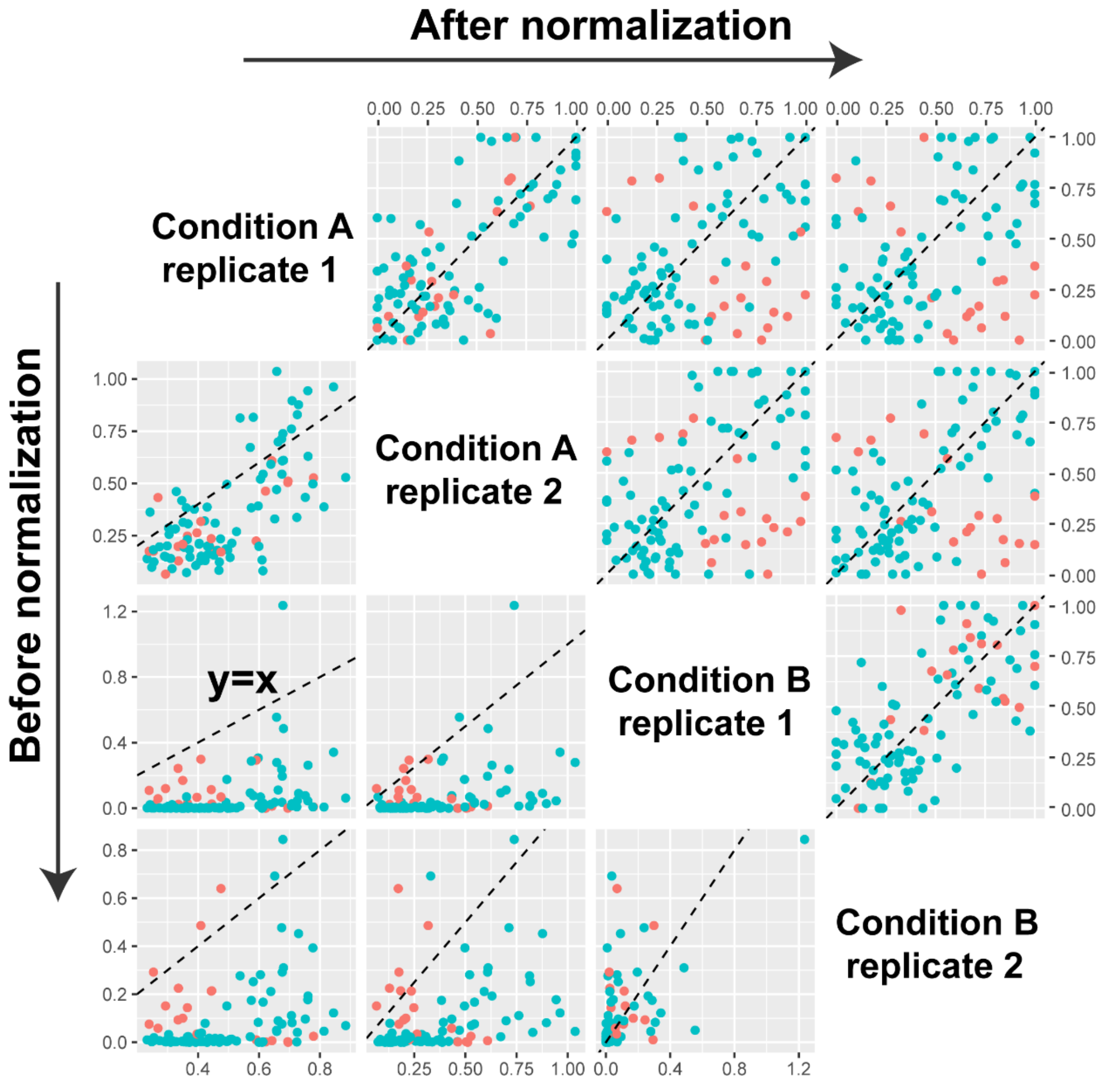


**Supplementary Figure 2: The normalization module of DiffScan.** This procedure removes systematic bias (lower panel) for reactivity replicates of condition A and B. Normalized reactivities are comparable across all replicates (upper panel).

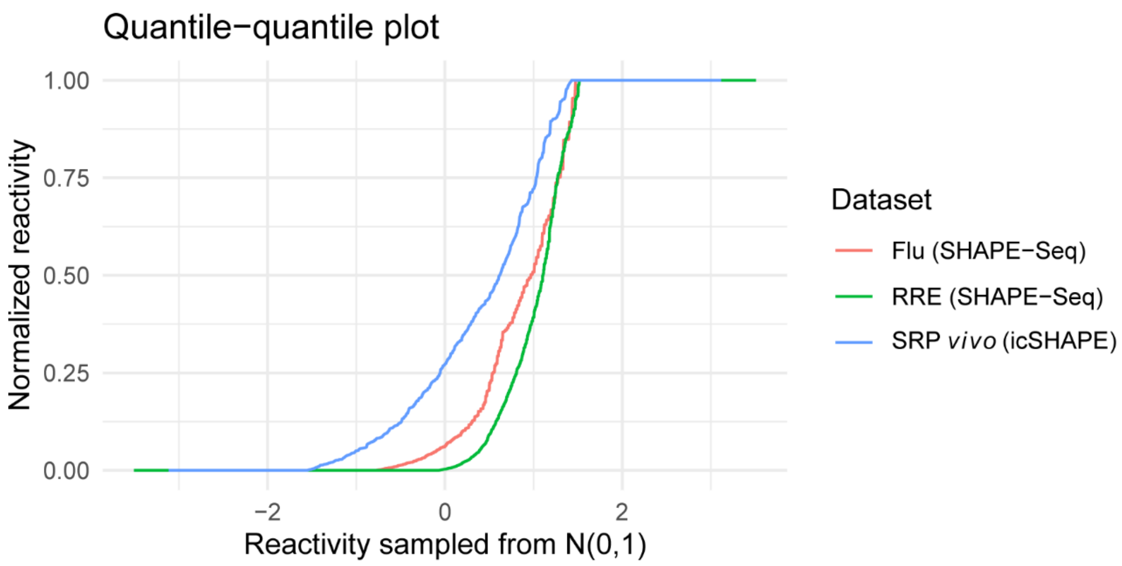


**Supplementary Figure 3: Quantile-quantile plot for normalized reactivity from the normalization module of DiffScan and reactivity sampled from N(0,1).** Normalized reactivities for three real datasets are displayed: the Flu and RRE dataset acquired using the SHAPE-Seq platform and the SRP *vivo* (*i.e.*, Control 6) dataset acquired using the icSHAPE platform (see Materials and Methods). Discrepancies of the distributions of normalized reactivities from normal distributions and the differences among the distributions of normalized reactivities demonstrate the necessity to take consideration of the diverse reactivity distributions from different SP platforms for subsequent differential analysis.

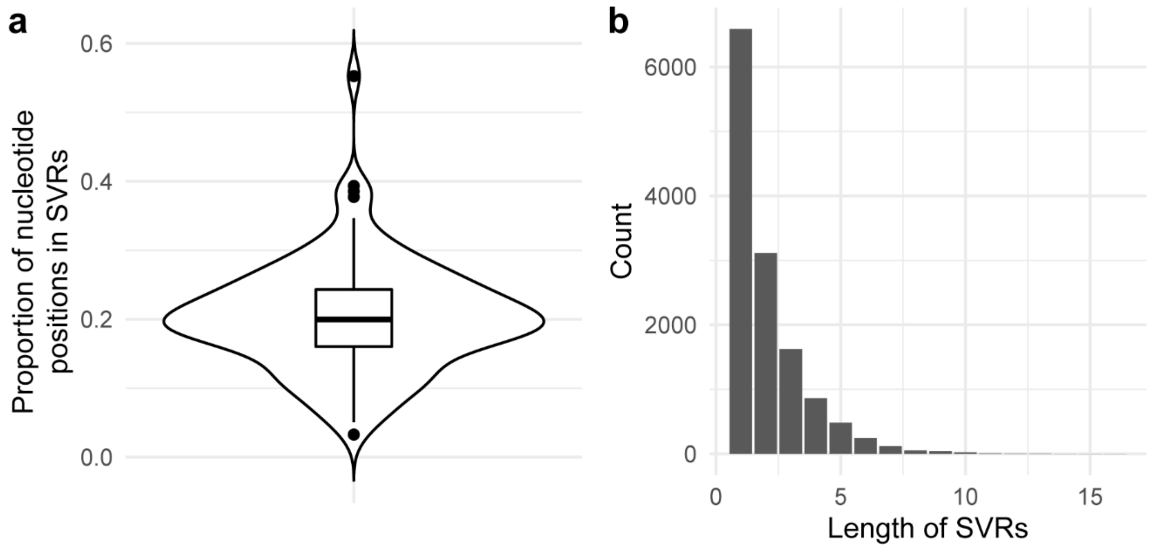


**Supplementary Figure 4: Details of the large-scale simulated datasets. a** Proportion of nucleotide positions that are in SVRs for each transcript. Boxplot elements: center line, median; box limits, upper and lower quartiles; whiskers, 1.5x interquartile range; points, outliers. **b** Length distribution of simulated SVRs.

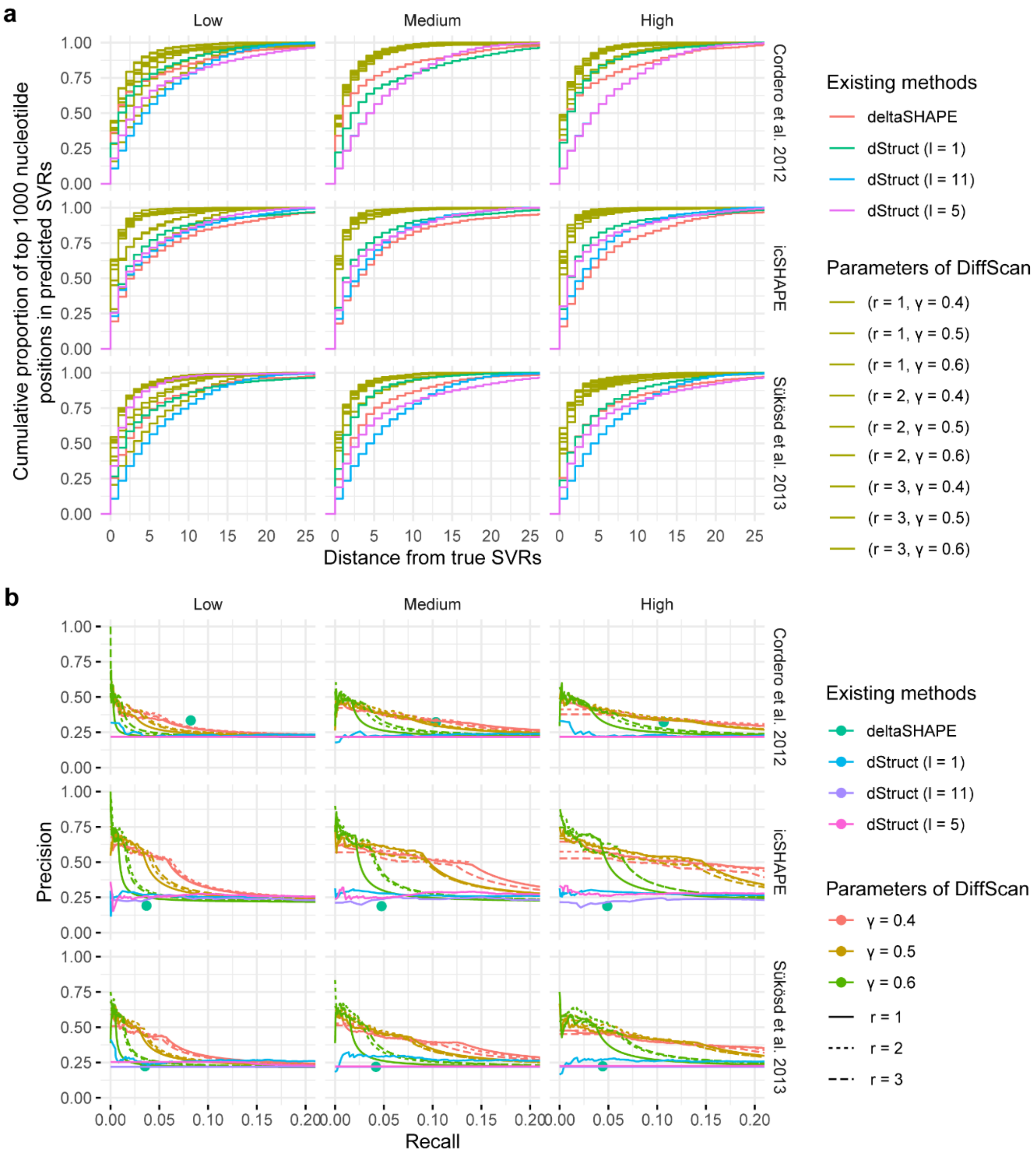


**Supplementary Figure 5: DiffScan maintains the relative advantage compared to dStruct and deltaSHAPE in the tested parameter settings.** For deltaSHAPE we used the default search length of 5 nt of the method; for dStruct we used search length 1 nt, 5 nt, and 11 nt. **a** Distances between simulated SVRs and top-ranked 1,000 nucleotide positions in predicted SVRs by existing methods and DiffScan with different parameters. **b** Precision-Recall curves for the prediction results from existing methods and DiffScan with different parameters. Rows: three types of reactivity models. Columns: three levels of strength of differential signals at simulated SVRs. Note deltaSHAPE does not allow external thresholding, and therefore it is represented as dots instead of curves.

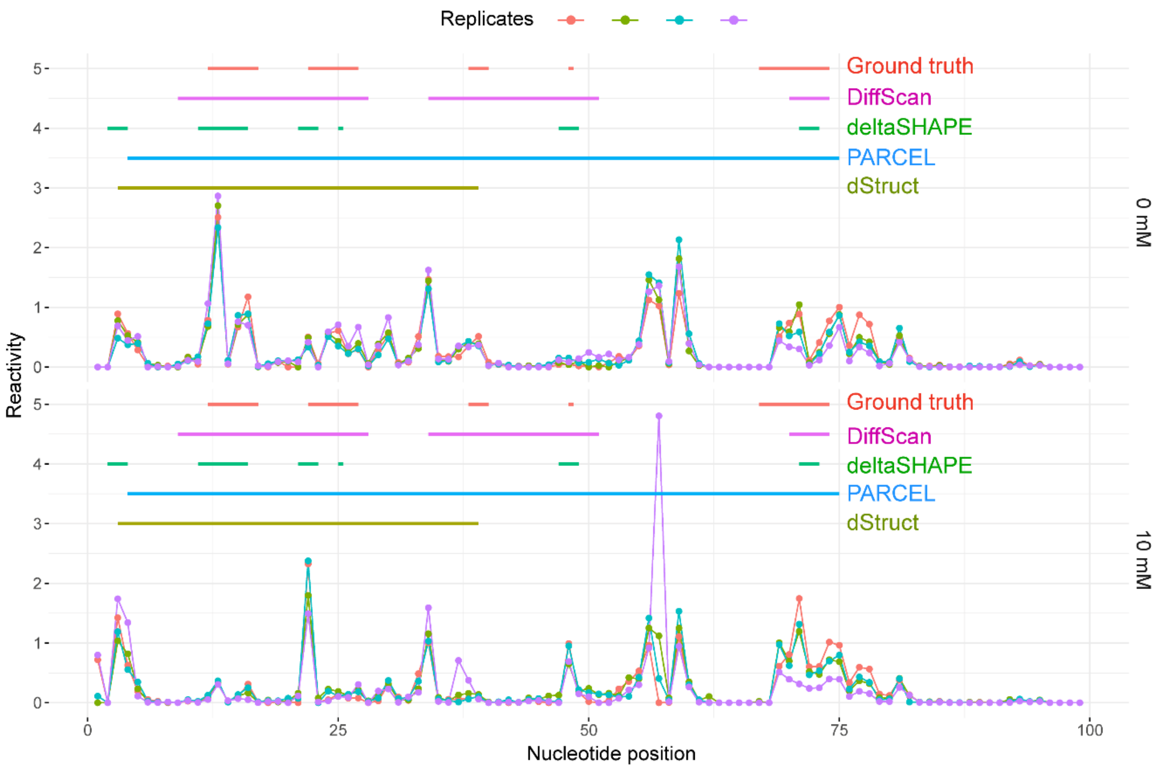


**Supplementary Figure 6: Predicted SVRs by different methods for the Flu dataset.** Top Panel: 4 reactivity replicates in the condition of 0 mM fluoride ions; bottom panel: 4 reactivity replicates in the condition of 10 mM fluoride ions. Line segments at the top denote true SVRs and predicted SVRs by different methods. RASA did not report any region.

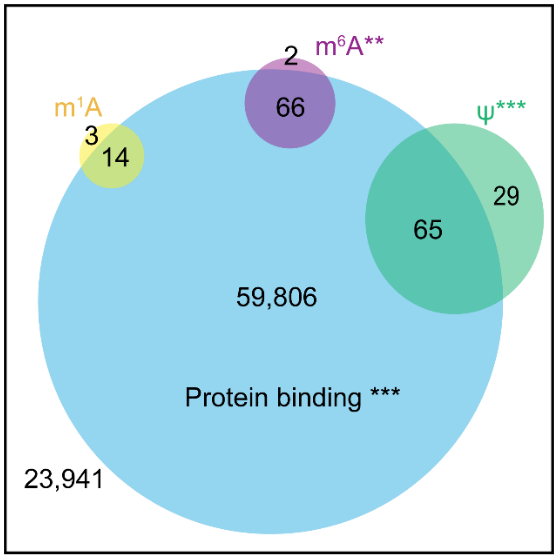


Np versus Cy
83,926 nucleotide positions in SVRs

**Supplementary Figure 7: Predicted SVRs by DiffScan for Np versus Cy were enriched with protein binding sites and RNA modification sites.** Np: nucleoplasm, Cy: cytoplasm. **p value (Fisher's exact test) < 1e-3, ***p value < 1e-6.

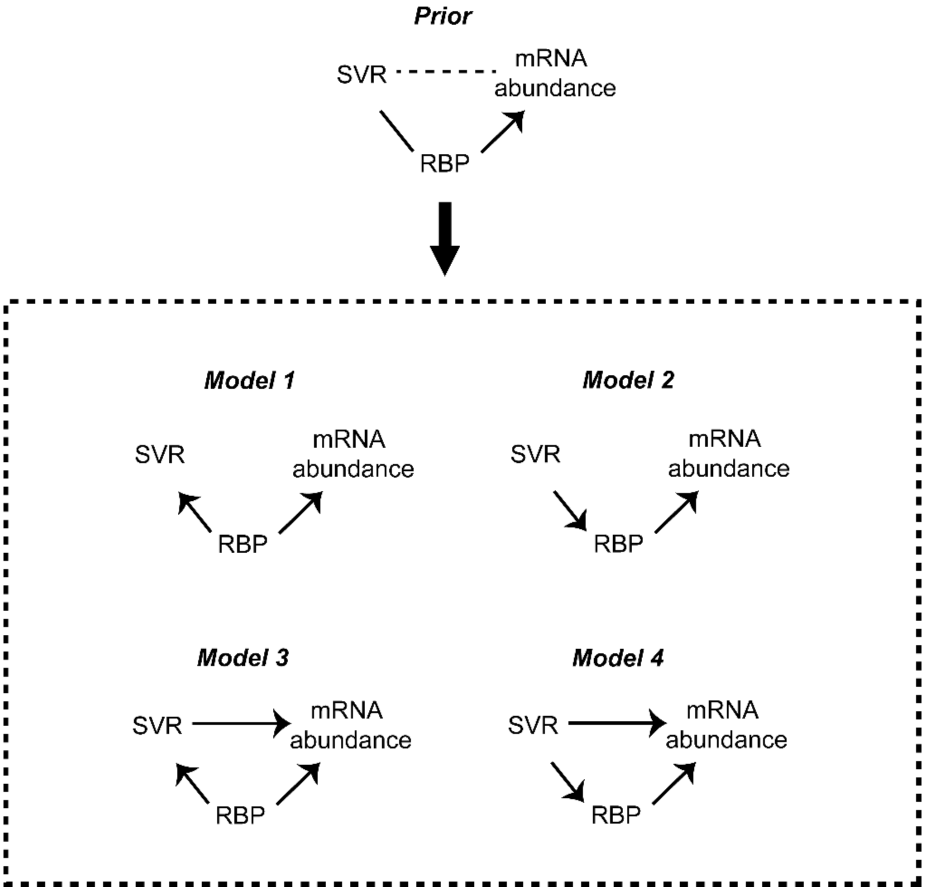


**Supplementary figure 8: Four models of which each describes the relationships among SVRs, changes in RBP binding, and changes in mRNA abundance.** Model 1-4 are based on the prior model in the top of the figure: the line connecting "SVR" and "RBP" in the figure stands for the detected enrichment for RBP in predicted SVRs. The arrow from "RBP" to "mRNA abundance" stands for the function of RBP in regulation of mRNA abundance, which is suggested for QKI and IGF2BP3 in literatures. The dotted line connecting "SVR" and "mRNA abundance" stands for the detected association between predicted SVRs and mRNA abundance; however, it is not clear whether SVRs regulate mRNA abundance.
